# A native prokaryotic voltage-dependent calcium channel with a novel selectivity filter sequence

**DOI:** 10.1101/783647

**Authors:** Takushi Shimomura, Yoshiki Yonekawa, Hitoshi Nagura, Michihiro Tateyama, Yoshinori Fujiyoshi, Katsumasa Irie

## Abstract

Voltage-dependent Ca^2+^ channels (Cavs) are indispensable for coupling action potentials with Ca^2+^ signaling in living organisms. The structure of Cavs is similar to that of voltage-dependent Na^+^ channels (Navs). It is known that prokaryotic Navs can obtain Ca^2+^ selectivity by negative charge mutations of the selectivity filter, but native prokaryotic Cavs had not yet been identified.

Here, we report the first identification of a native prokaryotic Cav, CavMr, and its relative, NavPp. Although CavMr contains a smaller number of negatively charged residues in the selectivity filter than artificial prokaryotic Cavs, CavMr exhibits high Ca^2+^ selectivity. In contrast, NavPp, which has similar filter sequence to artificial Cavs, mainly allows Na^+^ to permeate. Interestingly, a NavPp mutant whose selectivity filter was replaced with that of CavMr exhibits high Ca^2+^ selectivity. Mutational analyses revealed that the glycine residue of the CavMr selectivity filter is a determinant for Ca^2+^ selectivity. This glycine residue is well conserved among subdomains I and III of eukaryotic Cavs.

These findings provide new insight into the Ca^2+^ selectivity mechanism conserved from prokaryotes to eukaryotes.

## Introduction

Voltage-dependent Ca^2+^ channels (Cavs), which couple the membrane voltage with Ca^2+^ signaling, regulate some important physiological functions, such as neurotransmission and muscle contraction (Hille, 2001). The channel subunits of mammalian Cavs as well as mammalian voltage-dependent Na^+^ channels (Navs) have 24 transmembrane helices (24TM) (Catterall, 2000), and comprise 4 homologous subdomains with 6 transmembrane helices that correspond to one subunit of homo-tetrameric channels, such as Kvs and prokaryotic Navs (BacNavs). Comparison of the sequences between Navs and Cavs indicate that Navs derived from Cavs. Their two pairs of subdomains, domains I and III, and domains II and IV, are evolutionarily close to each other (Rahman et al., 2014; Strong et al., 1993). Therefore, the 24TM-type of Cavs are thought to have evolved from the single-domain type of Cavs with two times of domain duplications. Although prokaryotes are expected to have such ancestor-like channels, native prokaryotic Cavs have not yet been identified. The lack of ancestor-like prokaryotic Cavs is a missing link in the evolution of voltage-dependent cation channels.

In contrast to the lack of prokaryotic Cavs, many BacNavs have been characterized (Irie et al., 2010; Ito et al., 2004; Koishi et al., 2004; Lee et al., 2012; Nagura et al., 2010; Ren et al., 2001; Shimomura et al., 2016, 2011). The simple structure of BacNavs has helped to elucidate the molecular mechanisms of Nav (Irie et al., 2018, 2012; Tsai et al., 2013; Yue et al., 2002). In addition to this point, BacNavs has been used as the genetic tool for manipulating the neuronal excitability in vivo (Bando et al., 2014; Kamiya et al., 2019; Lin et al., 2010). The introduction of a several negatively charged amino acids into the selectivity filter of BacNavs leads to the acquisition of Ca^2+^ selectivity (Tang et al., 2013; Yue et al., 2002). Such a mutant of NavAb (a BacNav from *Arcobacter butzleri*) showed high Ca^2+^ selectivity, and the structural basis of Ca^2+^ selectivity has been discussed on the basis of its crystal structures (Tang et al., 2016, 2013). The selectivity filter sequences with a large number of aspartates in CavAb, which was made by mutations of NavAb, are quite different from those of the original mammalian Cavs.

Here, we newly characterized two BacNav homologues, CavMr from *Meiothermus ruber* and NavPp from *Plesiocystis pacifica*. These two channels are evolutionarily distant from the previously reported canonical BacNavs. We confirmed that CavMr has clear Ca^2+^ selectivity, and NavPp has Na^+^ selectivity with Ca^2+^-dependent inhibition. The discovery of these channels suggests the possible importance of voltage-regulated Ca^2+^ signaling in prokaryotes and may be a new genetic tool for controlling Ca^2+^ signaling. Furthermore, mutational analyses indicate that the glycine residue of the CavMr selectivity filter is important for Ca^2+^ selectivity. The glycine residue is also well conserved in the selectivity filter of the subdomain I and III of mammalian Cavs. On the basis of these observations, we propose that CavMr is an ancestral-type of native prokaryotic Cav with a Ca^2+^ selectivity mechanism different from that in artificial CavAb. CavMr and NavPp are expected to advance our understanding of Ca^2+^ recognition and the evolution of voltage-dependent cation channels.

## Results

### Identification of two prokaryotic channels with Ca^2+^ permeability and inhibition

We searched for the primary seqeunces of prokaryotic Cavs in the GenBank^TM^ database. In mammalian and prokaryotic Navs and Cavs, a larger number of negative charges in the filter increases Ca^2+^ selectivity (Heinemann et al., 1992; Tang et al., 2014; Yue et al., 2002). Several BLAST search rounds using the pore regions (S5-S6) of NaChBac (or NavBh; a BacNav from *Bacillus halodurans*) as templates revealed a series of candidate prokaryotic Cavs (Fig.1a) with a selectivity filter sequence similar to the “TLESW” motif, but more negatively charged: ZP_04038264 from *Meiothermus ruber*, ZP_01909854 from *Plesiocystis pacifica*, YP 003896792_from *Halomonas elongata*, and YP_003073405 from *Teredinibacter turnerae* (*SI Appendix*, Fig.S1a). These channels belong to a different branch of the phylogenic tree than that of canonical BacNavs (Fig.1a) and have some negatively charged residues in their selectivity filter, similar to CavAb (Fig.1b). In the case of the expression of prokaryotic channels, insect cells are better than mammalian cells to generate large current amplitudes (Irie et al., 2018). We therefore transfected Sf9 cells with these genes and measured the resulting whole-cell currents. Although the cells transfected with genes from *H. elongata* and *T. turnerae* failed to show detectable currents, those from *M. ruber* and *P. pacifica* showed currents in response to a depolarizing stimulus from a −140 mV holding potential (Fig.1c and d). To estimate the Ca^2+^ permeability, we measured their current-voltage relationships. The *M. ruber* channel had clearly larger currents in the high-Ca^2+^ solution than in the high-Na^+^ solution, and very positive reversal potential was observed in a high-Ca^2+^ bath solution (Fig.1e). In contrast, the currents derived from the *P. pacifica* channel increased with increases in the bath Na^+^ concentration and significantly decreased when the Na^+^ solution was replaced with a high Ca^2+^ solution. The reversal potential fit well to the Na^+^ equilibrium potential in the high-Na^+^ solution (Fig.1f). These current-voltage relationships suggest that the *M. ruber* channel has a preference for Ca^2+^ and the *P. pacifica* channel has a preference for Na^+^. Therefore, the two newly identified channels from *M. ruber* and *P. pacifica* are abbreviated as CavMr and NavPp respectively, based on their ion selectivity and species name.

**Figure 1.**
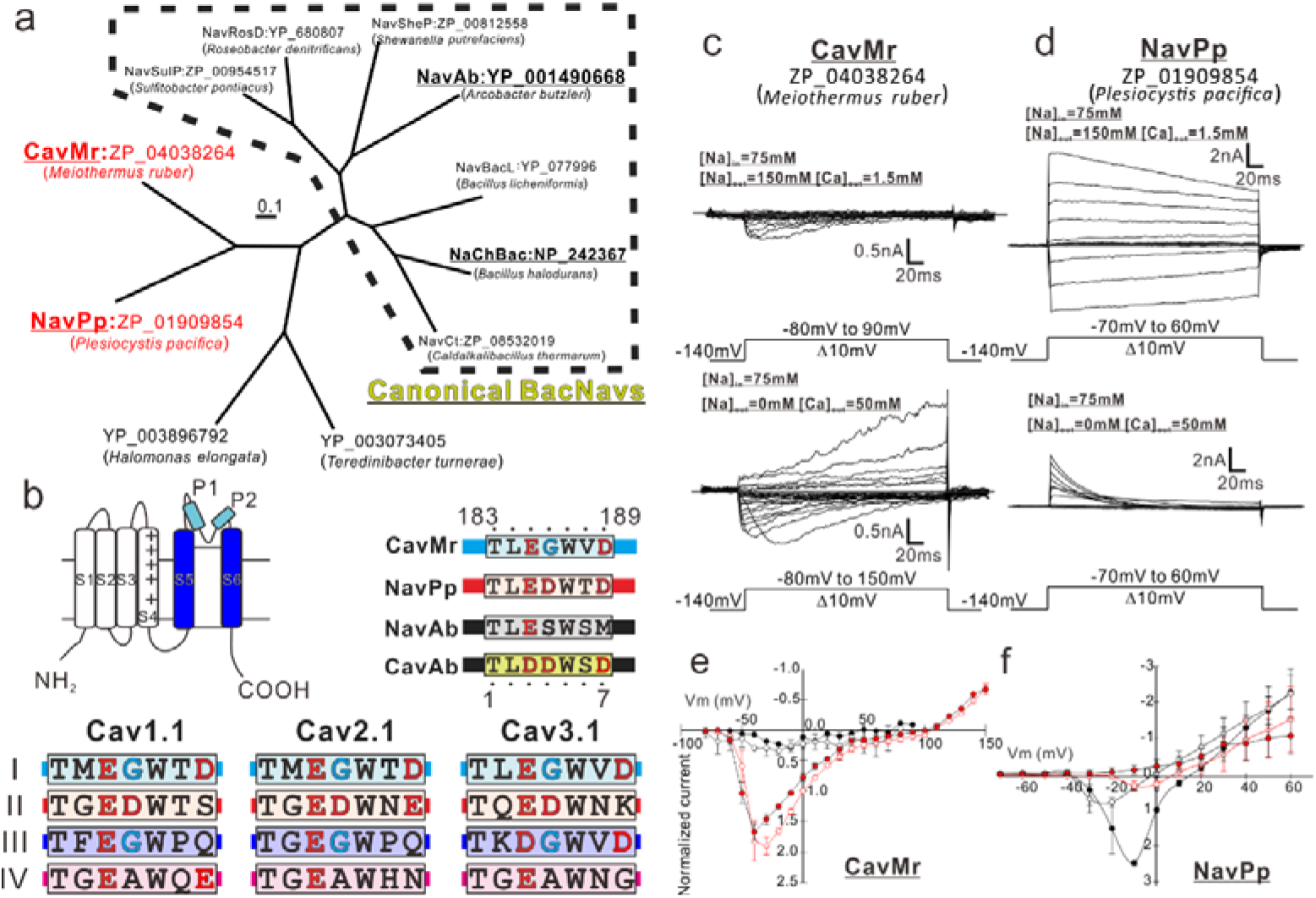
Sequence analysis and the representative current recordings of the novel BacNav homologues. a). Phylogenetic tree of the BacNav homologues with their GenBank^TM^ accession numbers. The ClustalW program was used to align the multiple protein sequences of the BacNav homologues. The phylogenetic tree was generated using “PROTDIST”, one of the PHYLIP package (Phylogeny Inference Package: http://evolution.genetics.washington.edu/phylip.html). The branch lengths are proportional to the sequence divergence, with the scale bar corresponding to 0.1 substitution per amino acid position. Four homologues colored that are not included in canonical BacNavs were cloned and expressed to check the current activity. Those of two which are underlined in red and shown as bold generated the detectable currents. b). Schematic secondary structure and selectivity filter sequence of BacNavs and human Cavs. Cylinder indicates α-helix. The selectivity filter sequences are indicated by alphabetical characters. Negatively charged residues are colored by red. Glycine residues in the position 4 are colored by cyan. The straight lines indicate the other part of pore domain. The selectivity filter sequence of hCav1.1 (UniProt ID: Q13698), hCav2.1 (O00555) and hCav3.1 (O43497), were used. c and d). Representative current traces to obtain the current-voltage relationships of CavMr (c) and NavPp (d) in Sf9 cells. The straight lines indicating the zero-current level in the representative current traces. Currents were generated under the bath solutions containing high Na^+^ (top) and high Ca^2+^ (middle), by a series of step-pulses shown in bottom. e and f). Current-voltage relationships of CavMr (e) and NavPp (f) measured under the different bath solutions (filled black; 150 mM NaCl, open black; 75 mM NaCl and 75 mM NMDG-HCl, open red; 75 mM NaCl and 50 mM CaCl_2_, filled red; 50 mM CaCl_2_ and 75 mM NMDG-HCl). Currents of CavMr and NavPp were normalized to that by 0mV depolarization stimuli under 75 mM NaCl and 50 mM CaCl_2_ bath solution and 150 mM NaCl bath solution, respectively.

To clearly compare the positions of the residues in the selectivity filter in each channel, we renumbered the seven residues comprising the selectivity filter in the following description. For example, the seven residues of the CavMr selectivity filter are 183-TLEGWVD-189, and thus Thr183 and Asp189 were renumbered to Thr1 and Asp7 (Fig.1b). Notably, the amino acid sequence of the selectivity filter in CavMr is similar to the conserved features of domains I/III in mammalian Cavs, a glycine at position 4 and a polar or negatively charged residue at position 7 (Fig.1b), which are not observed in the BacNav family. In addition, its sequence is quite similar to that of the human Cav subdomain I, or even the same as Cav3.1 and 3.2 (Fig.1b).

In the following experiments, to accurately evaluate the reversal potential for the ion selectivity analysis, we introduced a single mutation that resulted in large and long-lasting channel currents. T220A and G229A mutations in NaChBac led to slower inactivation, indicating suppression of the transition to the inactivated state (Irie et al., 2010; Shimomura et al., 2016). We introduced a T232A mutation to NavPp and a G240A mutation to CavMr, corresponding to the NaChBac mutations of T220A and G229A, respectively. These mutants stably showed measurable currents, even after they were administrated multiple depolarizing stimuli (*SI Appendix*, Fig.S1b-e).

### CavMr has high Ca^2+^ selectivity over Na^+^

We accurately quantified the selectivity of CavMr for Na^+^ and Ca^2+^ (*P*_Ca_/*P*_Na_) by measuring the reversal potential under bi-ionic conditions, in which the Ca^2+^ concentration in the bath solution was changed to 1.5, 4, 10, 20, and 40 mM while the intracellular Na^+^ concentration was held constant at 150 mM(Fig.2a and b). The plot of the reversal potentials as a function of [Ca^2+^] had a slope of 41.07±2.64 mV /decade (*n* = 4), a value close to that for Ca^2+^ (Fig. 2c), and indicated that CavMr had a *P*_Ca_/*P*_Na_ of 218 ± 38 (Fig.2a and b, Table1). This high *P*_Ca_/*P*_Na_ value is comparable to that of CavAb. Among several species of cations, including Sr^2+^, K^+^, and Cs^+^ (Fig.2d and e), Ca^2+^ had the highest permeability relative to Na^+^ (Fig.2f and g, Table1). On the basis of these results, CavMr was confirmed to be a native prokaryotic Cav with high Ca^2+^ selectivity. We also investigated whether CavMr shows the typical anomalous mole fraction effect (Almers and McCleskey, 1984) and the non-monotonic mole fraction effect observed in NaChBac (Finol-Urdaneta et al., 2014). CavMr did not allow Na^+^ permeation under Ca^2+^-free (0 mM CaCl_2_ and 1 mM EGTA) conditions (*SI Appendix*, Fig. S2a and b). Also, different from the recording of NaChBac currents in a solution containing Na^+^ and K^+^, CavMr had an apparently monotonic current increase depending on the Ca^2+^ mole fraction to Na^+^ (*SI Appendix*, Fig. S2c and d).

**Figure 2.**
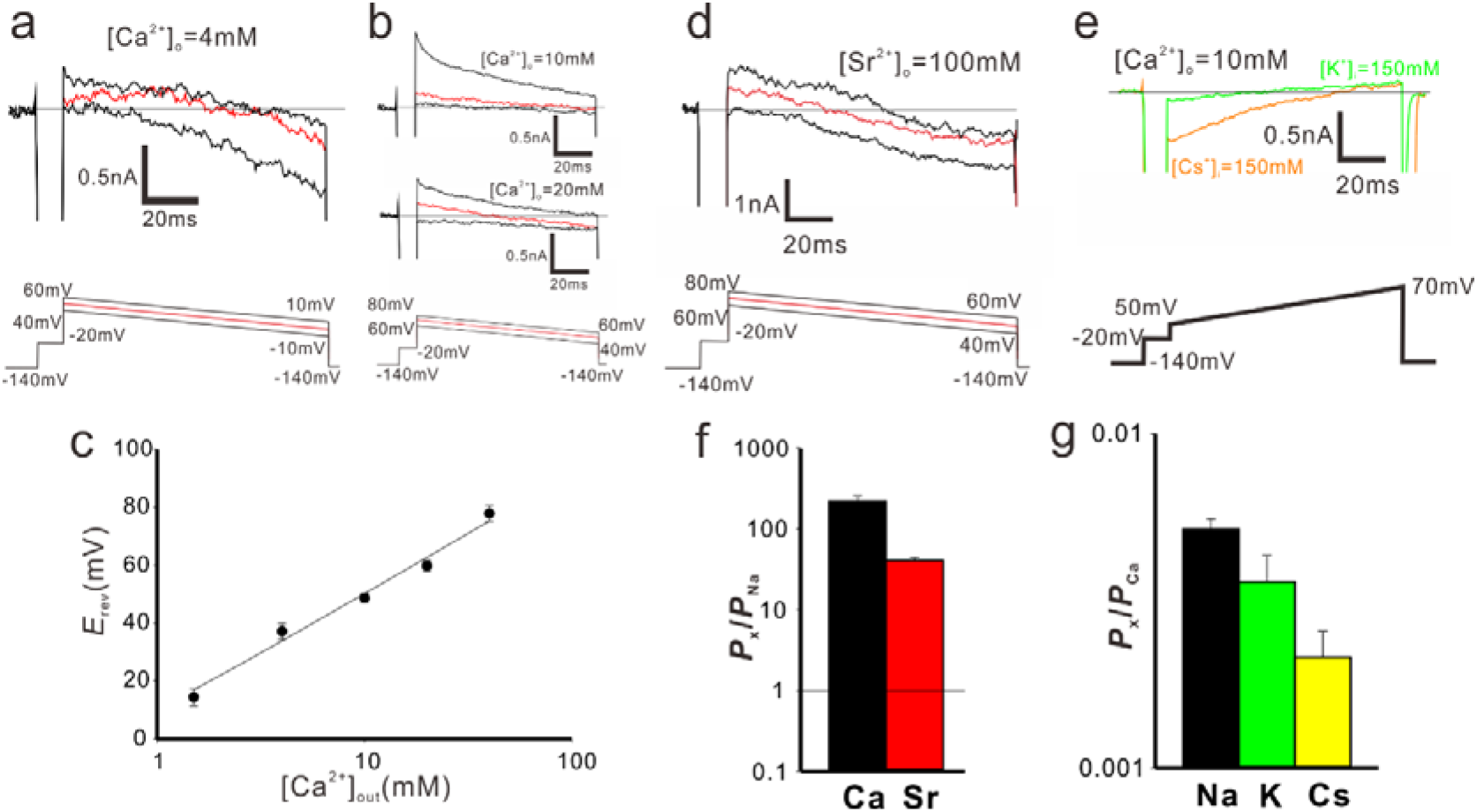
Cation selectivity of CavMr a and b). Recordings of the reversal potential of CavMr currents using the ramp protocol. Currents were generated by the step pulse of −20 mV from −140 mV holding potential, followed by the ramp pulses with different voltage values (shown at the bottom of panels a and b). The values of the reversal potential recorded with three different ramp pulses were averaged. Currents were measured under the bath solution containing 4 mM (a) and 10 and 20 mM (b) CaCl_2_ and the pipette solution with 150 mM NaCl. c). The plot of the reversal potential to the bath [Ca^2+^]_out_. Each value was obtained using the protocol shown in a and b. The relationship was fitted by a line with the slope of 41.07±2.64 mV per decade (*n* = 4). d). Representative current traces to obtain the reversal potential under the condition of 100 mM [Sr^2+^]_out_ and 150 mM [Na^+^]_in_. Currents were generated by the protocol shown in the lower part. e). Representative current traces to investigate the *P*_Cs_/*P*_Ca_ and *P*_K_/*P*_Ca_, the pipette solutions contained 150 mM [Cs^+^]_in_ for *P*_Cs_/*P*_Ca_ and 150 mM [K^+^]_in_ for *P*_K_/*P*_Ca_, while the bath solution contained 10 mM [Ca^2+^]_out_ in both cases. f). The relative permeability of Ca^2+^ or Sr^2+^ to Na^+^ in CavMr, calculated from the reversal potential that was obtained in a, b, and d. g). The relative permeability of each monovalent cation to Ca^2+^ in CavMr, derived from the data shown in e.

**Table 1.** Relative permeability of CavMr and NavPp

| | $P_{\text{Ca}}/P_{\text{Na}}$ | $P_{\text{Sr}}/P_{\text{Na}}$ | $P_{\text{K}}/P_{\text{Na}}$ | $P_{\text{Cs}}/P_{\text{Na}}$ |
| --- | --- | --- | --- | --- |
| CavMr G240A | 218 ± 38<br>($n= 20$ ) | 40.6 ± 3.4<br>($n= 6$ ) | 0.0036±0.00072<sup>a</sup><br>($n= 4$ ) | 0.0021±0.00042<sup>b</sup><br>($n= 4$ ) |
| Pp | 13.8 ± 2.0<br>($n= 7$ ) | 24.5 ± 0.3<br>($n= 5$ ) | 0.95 ± 0.04<br>($n= 4$ ) | 0.57 ± 0.05<br>($n= 3$ ) |
| G4D | 7.73 ± 2.24<br>($n= 11$ ) | 18.6 ± 6.1<br>($n= 4$ ) | 1.20 ± 0.28<br>($n= 4$ ) | 0.87 ± 0.21<br>($n= 4$ ) |
| G4S | 11.9 ± 1.5<br>($n= 5$ ) | 4.23 ± 0.27<br>($n= 5$ ) | 1.54 ± 0.12<br>($n= 5$ ) | 2.02 ± 0.48<br>($n= 3$ ) |
| V6T | 40.1 ± 9.7<br>($n= 5$ ) | 13.3 ± 2.5<br>($n= 5$ ) | 0.69 ± 0.26<br>($n= 3$ ) | 0.54 ± 0.60<br>($n= 3$ ) |
| D7M | 144 ± 12<br>($n= 5$ ) | 20.7 ± 2.7<br>($n= 5$ ) | N.D. | N.D. |
| NavPp T232A | 0.308±0.028<br>($n= 10$ ) | 0.38±0.027<br>($n= 9$ ) | 0.16±0.026<br>($n= 9$ ) | 0.0052±0.0006<br>($n= 7$ ) |
| Mr | 215 ± 33<br>($n= 7$ ) | 86.3 ± 12.2<br>($n= 4$ ) | 0.0045±0.00072<sup>a</sup><br>($n= 4$ ) | 0.0135±0.0039<sup>b</sup><br>($n= 9$ ) |
| D4G | 41.4 ± 6.7<br>($n= 10$ ) | 8.85 ± 0.95<br>($n= 4$ ) | << 0.01<br>($n= 3$ ) | << 0.01<br>($n= 4$ ) |
| D4S | 1.49 ± 0.18<br>($n= 6$ ) | 0.47 ± 0.08<br>($n= 6$ ) | 0.24 ± 0.01<br>($n= 4$ ) | 0.11 ± 0.02<br>($n= 3$ ) |
| T6V | 1.72 ± 0.19<br>($n= 10$ ) | 33.9 ± 5.0<br>($n= 8$ ) | 0.99 ± 0.03<br>($n= 4$ ) | 0.84 ± 0.02<br>($n= 4$ ) |

<sup>a</sup> Because of high $\text{Ca}^{2+}$ selectivity, $P_{\text{K}}/P_{\text{Ca}}$ were indicated
<sup>b</sup> Because of high $\text{Ca}^{2+}$ selectivity, $P_{\text{Cs}}/P_{\text{Ca}}$ were indicated

Studies of an artificial Cav, CavAb, revealed that Ca^2+^ selectivity depends on a large number of aspartates in the filter sequence (Tang et al., 2013). The high Ca^2+^ selectivity in CavMr was unexpected because the filter sequence contained only one aspartate residue (Fig.1b). Furthermore, CavMr-D7M, which has only one negatively charged residue in the selectivity filter “TLEGWVM”, still had high Ca^2+^ selectivity comparable to that of wild-type CavMr (*P*_Ca_/*P*_Na_ = 144 ± 12;*SI Appendix*, Fig. S2e and f and Table1). These findings indicate that CavMr and artificial CavAb have different Ca^2+^ selective mechanisms.

### NavPp is permeable to Na^+^ and is blocked by extracellular Ca^2+^

The current-voltage relationships of NavPp revealed a preference for Na^+^ (Fig.1d and f). Interestingly, NavPp, although having one more aspartate in the selectivity filter than CavMr, exhibited larger currents in Na^+^ solutions than in Ca^2+^ solutions (Fig.1b and d). Recordings in bath solution containing both Na^+^ and Ca^2+^ demonstrated that increasing the extracellular Ca^2+^ decreased the current in NavPp and led to a positive shift in the voltage dependence, indicating that a higher concentration of Ca^2+^ inhibited NavPp (Fig.1f and Fig. 3a and *SI Appendix*, Fig. S3a). With NavAb, increasing the Ca^2+^ concentration with a constant Na^+^ concentration in the bath solution led to a small increase in the current amplitude, probably due in part to Ca^2+^ permeability (Fig. 3a and *SI Appendix*, Fig. S3b). We also investigated the dependence of the direction of current flow on Ca^2+^ inhibition by comparing pipette solutions containing 10 mM or 150 mM Na^+^, in which the current flowed in an inward or outward direction, respectively, even under the same −10‐ mV depolarizing stimulus (Fig. 3a and *SI Appendix*, Fig. S3a and c). The results demonstrated that the inhibitory effects of Ca^2+^ on NavPp were independent of the current direction.

**Figure 3.**
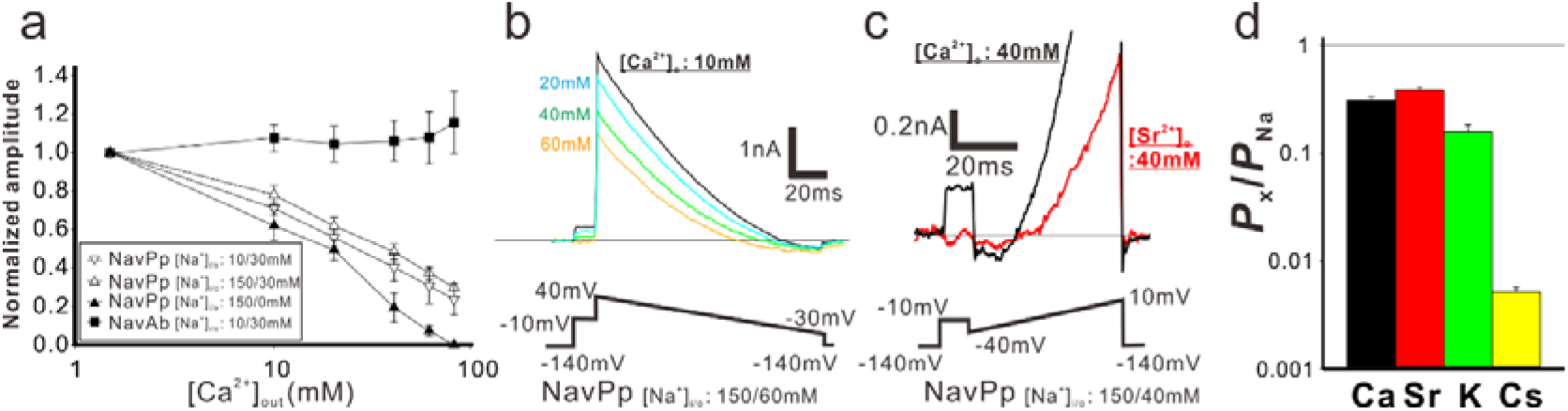
Na^+^ permeability and the extracellular Ca^2+^-dependent inhibition in NavPp. a) [Ca^2+^]_out_ dependent inhibitory effects in NavPp. Currents were normalized to those under 1.5 mM [Ca^2+^]_out_(filled square; NavAb current in the 30 mM [Na^+^]_out_ and 10 mM [Na^+^]_in_ (representative data are shown in Fig. S3b), open triangle down; NavPp inward current in the 30 mM [Na^+^]_out_ and 10 mM [Na^+^]_in_ (Fig. S3a), open triangle up; NavPp outward current in the 30 mM [Na^+^]_out_ and 150 mM [Na^+^]_in_ (Fig. S3c), filled triangle up; NavPp outward current in the 0 mM [Na^+^]_out_ and 150 mM [Na^+^]_in_. b). Representative currents to obtain the reversal potential in NavPp for *P*_Ca_/*P*_Na_. Currents were generated by the ramp protocol shown in bottom. [Ca^2+^]_out_ was varied from 10 to 60 mM with the fixed [Na^+^] both in the bath (60mM) and pipette (150mM). c). Representative currents to obtain the reversal potential in NavPp for *P*_Sr_/*P*_Na_. Currents were generated by the ramp protocol shown in bottom. [Sr^2+^]_out_ was 40 mM with the fixed [Na^+^] both in the bath (40mM) and pipette (150mM). d). The relative permeability of different cation species to Na^+^ in NavPp, calculated from the reversal potential that was obtained in b, c and Fig.S3d and e.

We then compared the relative permeability of various cations with that of Na^+^ in NavPp. In bi-ionic conditions with high concentration of Ca^2+^ in the bath and high Na^+^ in the pipette, the outward current was completely blocked by Ca^2+^ (Fig. 3a), and the inward current was hardly observed. The reversal potential was obtained under an extracellular solution containing Na^+^ ions, however, despite a partial Ca^2+^ or Sr^2+^-induced block (Fig. 3b and c). The selectivity of NavPp was higher for Na^+^ than for Ca^2+^, Sr^2+^, K^+^, and Cs^+^ (Fig. 3d and *SI Appendix*, Fig. S3d and e). The *P*_Ca_/*P*_Na_ was 0.308 ± 0.028 in a bath solution containing both Ca^2+^ and Na^+^, suggesting that a larger fraction of Ca^2+^ is allowed to permeate with outside Na^+^ ions through NavPp than through canonical BacNavs. Similar to Ca^2+^, Sr^2+^ also blocked the NavPp current, but may also permeate the channel along with Na^+^ ions (Fig. 3c). These findings demonstrate a unique feature of NavPp, a low affinity Ca^2+^ block, which is not reported in canonical BacNavs.

Interestingly, the filter sequence of NavPp, "TLEDWTD", has three negatively-charged residues, similar to the filter sequences of the artificial Ca^2+^-selective BacNav mutants (“TLEDWSD” mutant of NavAb and “TLEDWAD” mutant of NaChBac) (Tang et al., 2013; Yue et al., 2002). NavPp does not show Ca^2+^ permeability, however, but rather a Ca^2+^ block. We also investigated NavPp mutants with the same filter sequences as the artificial Cavs. NavPp-T6S “TLEDWSD” exhibited Ca^2+^-blocked currents similar to wild-type NavPp (*SI Appendix*, Fig. S3f). Further, NavPp-T6A “TLEDWAD” showed no inward current in bath solutions containing divalent cations, suggesting that the Ca^2+^-induced block was enforced. Therefore, both of the selectivity filter sequences providing Ca^2+^ selectivity to canonical BacNavs failed to generate Ca^2+^-permeable NavPp, indicating that the cation permeable mechanism of NavPp differs from that of canonical BacNavs, as well as that of CavMr.

### Swapping the filter regions between CavMr and NavPp revealed the importance of the glycine residue at position 4 for Ca^2+^ selective permeation

To search for the determinants of Ca^2+^ selectivity in CavMr, we investigated a series of mutants in which the filter regions were swapped between CavMr and NavPp (Fig.4a and b). The mutants with filter sequences swapped between NavPp and CavMr exhibited channel activity (*SI Appendix*, Fig. S4). A NavPp mutant whose selectivity filter was replaced with that of CavMr, named NavPp-Mr, exhibited much higher Ca^2+^ selectivity (*P*_Ca_/*P*_Na_ = 215 ± 33) as well as high Sr^2+^ selectivity, comparable to that of CavMr (Fig. 4c). In addition, NavPp-Mr excluded Cs^+^ similar to CavMr, but weakly allowed K^+^ permeation in contrast to CavMr. On the other hand, a CavMr mutant whose selectivity filter was replaced with that of NavPp (CavMr-Pp) almost lost its Ca^2+^ selectivity (*P*_Ca_/*P*_Na_ = 13.8 ± 2.0), and was less able to discriminate Cs^+^ and K^+^ from Na^+^ (Fig. 4d). That is, CavMr-Pp was a more non-selective channel than the wild-type CavMr, rather than a Na^+^-selective channel. Namely, the Ca^2+^ selectivity (from NavPp to CavMr) was almost transferable, but the Na^+^ selectivity was not. We also investigated the full swapping of the filter sequences between CavMr and NavAb (Fig. 4a), but neither swapped mutant of CavMr and NavAb had detectable currents. This finding suggested that CavMr and NavAb achieve cation selectivity by different structural backbones and mechanisms.

**Figure 4.**
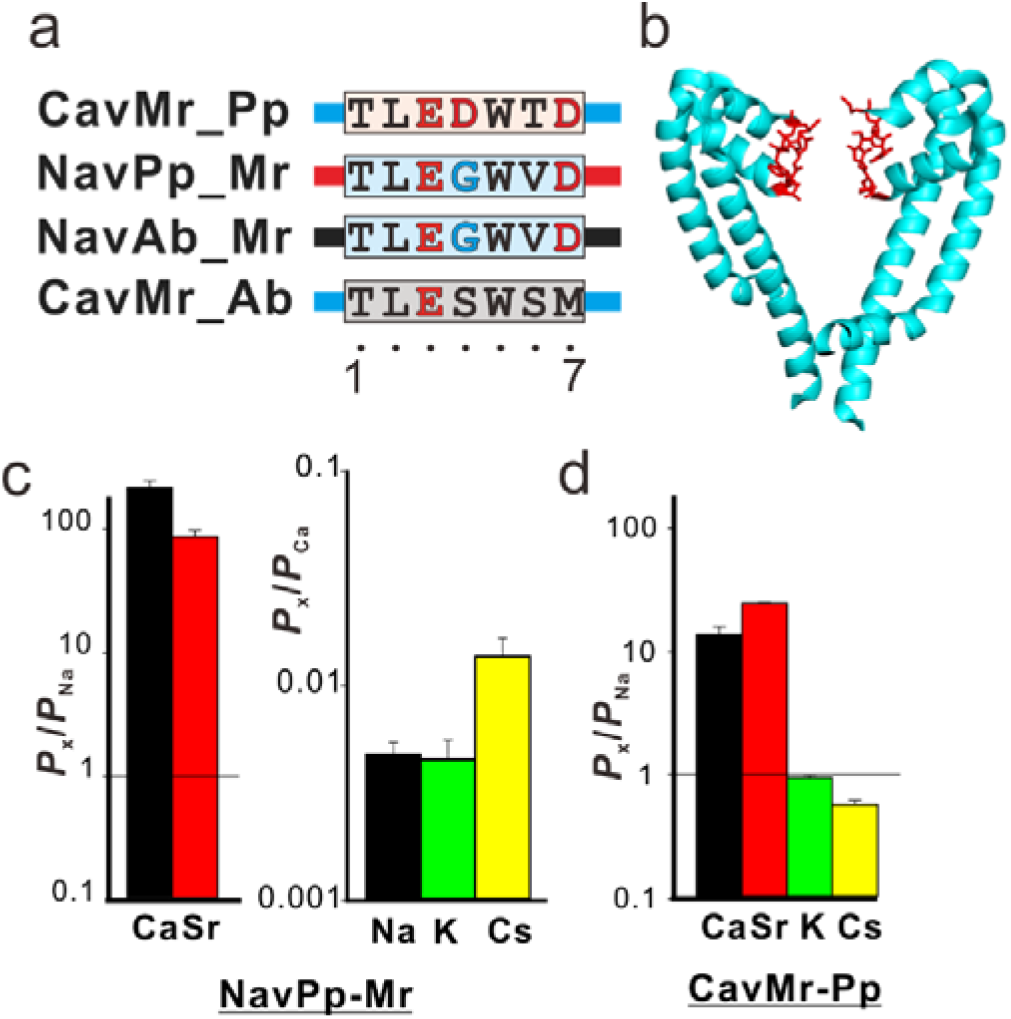
The cation selectivity of the swapping mutant channels in their selectivity filter between CavMr and NavPp. a) Amino acid sequences of the selectivity filter in the swapped mutants, CavMr-Pp, Nav-Pp, NavAb_Mr, and CavMr_Ab. The selectivity filter sequences of CavMr, and NavPp, and NavAb are indicated by alphabetical characters with cyan, red, and gray shade, respectively. Negatively charged residues are colored by red. Glycine residues are colored by cyan. The straight lines of cyan, red, and black indicates the other part of pore domain of CavMr, NavPp, and NavAb, respectively. b) Pore domains of crystal structure of NavAb (PDB code:5YUA). The selectivity filter, which corresponds to the sequences shown in a, was indicated in red. c) The relative permeability of divalent cations to Na^+^ (left) and that of monovalent cations to Ca^2+^ (right) in NavPp-Mr. d) The relative permeability of different cation species to Na^+^ in CavMr-Pp.

Positions 4 and/or 6 of the filter sequences are thought to be important for Ca^2+^-selective permeation in NavPp-Mr and CavMr, because only these two positions were mutated in the swapping experiments. We investigated which of the mutations in positions 4 and 6 had greater effects on the loss of and acquisition of Ca^2+^ selectivity in CavMr and NavPp, respectively. In CavMr, both of two single mutants, CavMr-G4D and CavMr-V6T, decreased Ca^2+^ selectivity and allowed K^+^ and Cs^+^ permeation (Fig. 5a, *SI Appendix*, Fig.S5). Especially, the mutational effect was greater in CavMr-G4D, whose *P*_Ca_/*P*_Na_ was less than 10 (7.73 ± 2.24). CavMr-G4S, in which Gly4 was replaced with the Ser4 of NavAb, also exhibited lower Ca^2+^ selectivity (*P*_Ca_/*P*_Na_ = 11.9 ± 1.5) and was also K^+^ and Cs^+^ permeable, indicating that even a minor substitution by serine is not tolerable and does not allow for the selection of specific cations (Fig. 5b, Table1, and *SI Appendix*, Fig. S6). In the case of NavPp, NavPp-D4G acquired Ca^2+^ selectivity over Na^+^, and also showed a greater exclusion to K^+^ and Cs^+^ than wild-type NavPp (Fig. 5c and *SI Appendix*, Fig.S5). In contrast, NavPp-T6V failed to acquire the high Ca^2+^ selectivity (*P*_Ca_/*P*_Na_ = 1.72 ± 1.09) and also allowed K^+^ and Cs^+^ permeation, while it had relatively high Sr^2+^ selectivity. These results indicate that, in both CavMr and NavPp, a glycine residue at position 4 is a key determinant for Ca^2+^ selectivity. It is noteworthy that the glycine is a conserved residue at position 4 of subdomains I and III in all subtypes of mammalian Cavs (Fig.1b).

**Figure 5.**
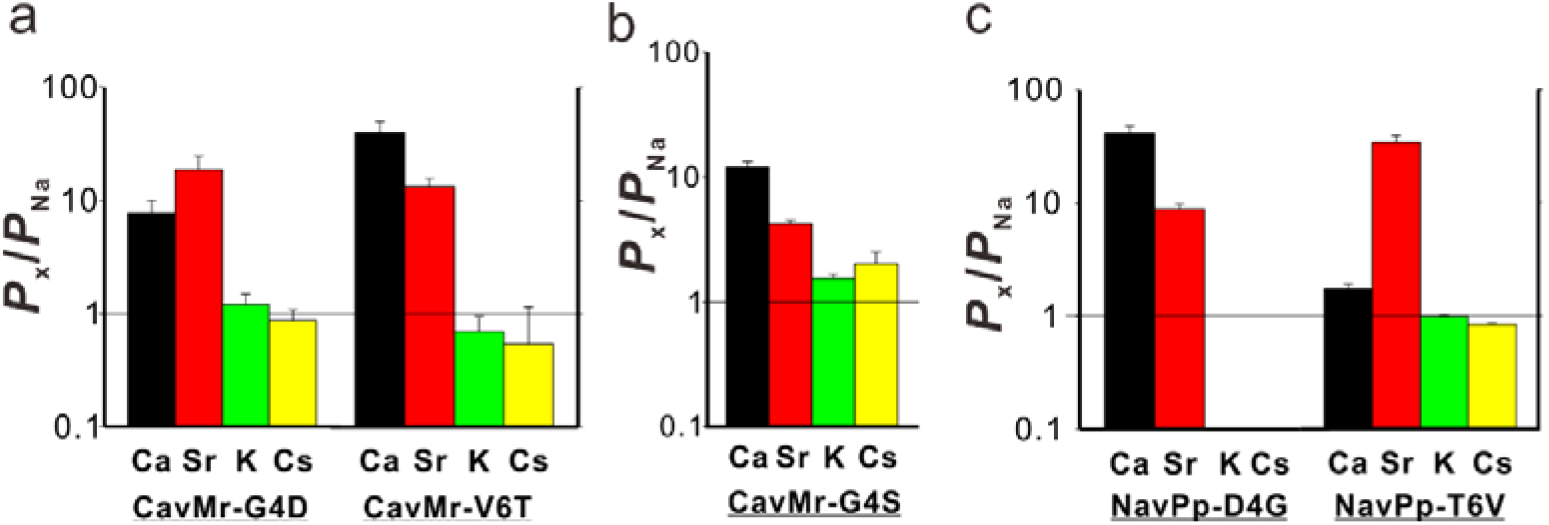
The single point mutations losing and obtaining Ca^2+^ selectivity of CavMr and NavPp, respectively. a) The relative permeability of each cation species to Na^+^ in the single-point mutants of CavMr. The selectivity filter of CavMr was changed to the corresponding residues of NavPp at position 4 (G4D) and position 6 (V6T), respectively b) The relative permeability of each cation species to Na^+^ in the G4S mutants of CavMr, whose position 4 residue of the selectivity filter was mutated to the corresponding residue of canonical BacNavs. c) The relative permeability of each cation species to Na^+^ in the single-point mutants of NavPp. The selectivity filter of NavPp was changed to the corresponding residues of CavMr at position 4 (D4G) and position 6 (T6V), respectively.

## DISCUSSION

### A native prokaryotic voltage-dependent Ca^2+^ channel has a unique Ca^2+^ selective mechanism

In this study, we newly characterized two prokaryotic voltage-dependent cation channels, CavMr and NavPp. CavMr is the first native prokaryotic Cavs reported despite its BacNav-like “TxExW” motif, and NavPp could be inhibited by high concentrations of extracellular Ca^2+^. The *P*_Ca_/*P*_Na_ of CavMr was more than 200 (Fig. 2e and Table 1), comparable to that of CavAb, an artificial Ca^2+^ channel. Anomalous mole fraction effects were not observed in CavMr (*SI Appendix*, Fig. S2a and b), suggesting that CavMr has a very high affinity for Ca^2+^. In addition to providing new insights about general Ca^2+^-selective mechanisms, CavMr has the potential to be a new genetic tool for upregulating calcium signaling, as BacNavs are useful genetic tools for increasing action potential firing in mice (Bando et al., 2014; Kamiya et al., 2019; Lin et al., 2010). Phylogenetic analysis demonstrated that CavMr and NavPp are similar to each other, but distant from canonical BacNavs (Fig. 1a). The high Ca^2+^ selectivity of CavMr was transferable to NavPp. Intriguingly, two pairs of mutants with the same selectivity filter (CavMr-G4D and NavPp-T6V, CavMr-V6T and NavPp-D4G) showed a very similar tendency with regard to both the order and extent of cation selectivity (Fig. 5a and c). Therefore, the basic overall architecture of the NavPp selectivity filter could be similar to that of CavMr. On the other hand, the Ca^2+^ selectivity mechanism of CavMr completely differs from that of CavAb. Structural comparison of NavAb and CavAb showed that the aspartate mutations did not alter the main chain trace, and simply introduced the negative charges around the ion pathway to increase Ca^2+^ permeability (Fig. 7a and b) (Tang et al., 2013). In contrast, in the case of CavMr, two non-charged residues (Gly4 and Val6) are required for the high Ca^2+^ selectivity (Fig. 4c, 5a), but Asp7 is not necessary (*SI Appendix*, Fig. S2f). A no-charge mutation at position 7, CavMr-D7M “TLEGWVM”, is an outstanding example demonstrating that high Ca^2+^ selectivity can be achieved in the absence of any aspartates in its filter region (*SI Appendix*, Fig. S2f). Furthermore, the introduction of a negative charge into the selectivity filter (G4D mutation) had the opposite effect on the Ca^2+^ selectivity of CavMr compared with NavAb and NaChBac (Tang et al., 2014; Yue et al., 2002). Moreover, the decreased selectivity in G4S also indicates that the glycine at position 4 is indispensable for Ca^2+^ selectivity in CavMr (Fig. 5b). The flexibility and/or small size of the glycine at position 4 in CavMr might be critical. These findings are inconsistent with the notion derived from the Ca^2+^-selective mutants of NavAb and NaChBac, and therefore the native structure of the selectivity filter and the molecular mechanism of ion selectivity of CavMr are thought to differ from those of CavAb. While the structure of CavMr is not yet available, we are able to speculate on the structure of the selectivity filter of CavMr on the basis of the structure of human Cav1.1 subdomains I and III (Wu et al., 2016) (Fig. 6c), whose selectivity filter sequences are very similar to that of CavMr. In the selectivity filter of Cav1.1 subdomains I and III, the side chain of the residue at position 7 is shifted outward. The position-4 glycine residue widens the entrance of the selectivity filter, which would facilitate the entry of hydrated cations into the ion pore and might increase Ca^2+^ selectivity.

**Figure 6.**
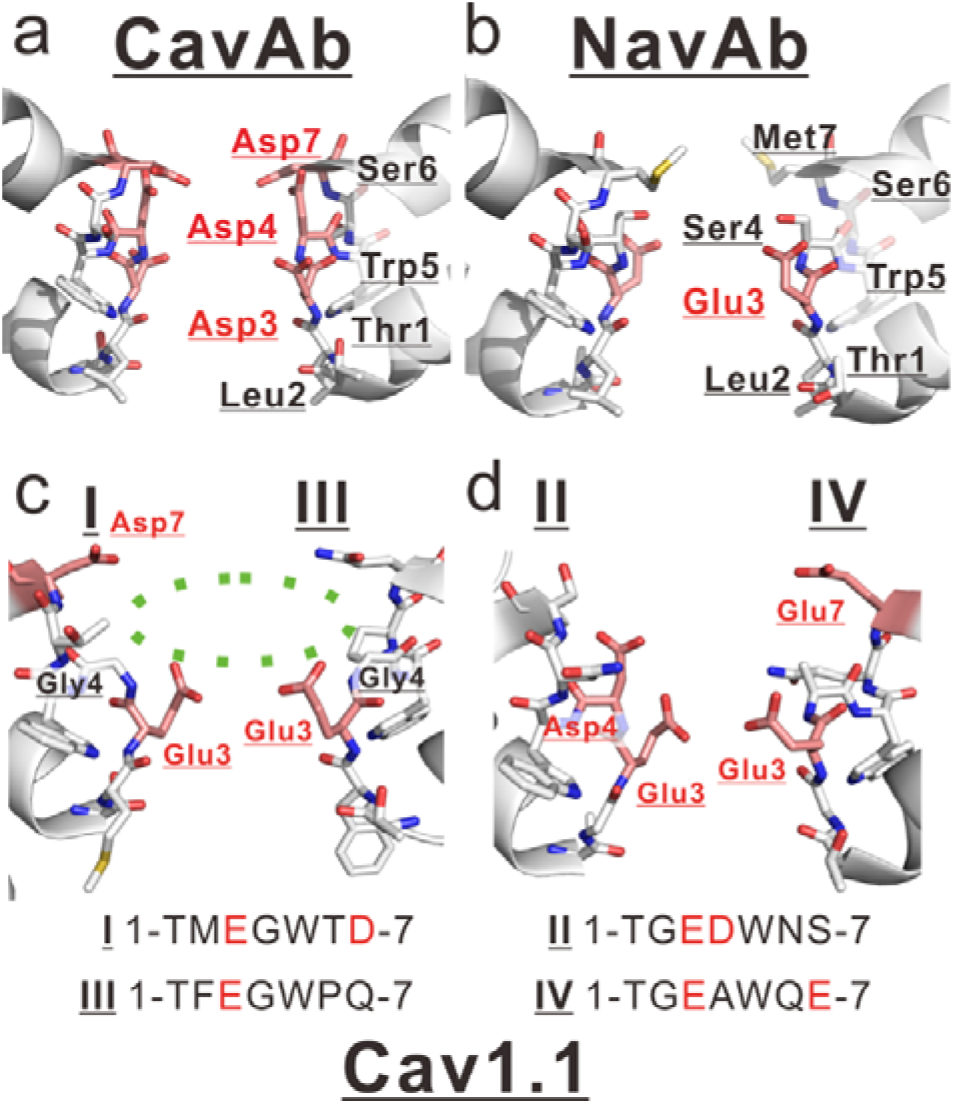
Comparison between mammalian and prokaryotic Cav. a and b). Structures of the selectivity filter in CavAb (PDB code: 4MVZ) and NavAb (PDB code: 5YUA). c and d) Structure of the rabbit Cav1.1 selectivity filter (PDB code: 5GJV). The subdomains I and III (c), and II and IV (d) were separately shown. The carbon atoms of negatively charged residues were indicated in pink. Dashed green circle indicates the wide entrance of the selectivity filter.

### Cavs in prokaryotes and the species-specific tuning of homo-tetrameric channels

Prokaryotes have a number of putative Ca^2+^ binding proteins, such as EF-hand proteins, P-type Ca^2+^ pumps, and Ca^2+^ transporters (Domínguez et al., 2015). The intracellular Ca^2+^ concentration is kept low and changes in response to mechanical and chemical stimuli (Dominguez, 2004). These features imply that prokaryotic Ca^2+^ signaling is similar to that of eukaryotes. The strong ability of CavMr to exclude Na^+^ and K^+^ along with Ca^2+^ permeation suggests that its primary physiological role is Ca^2+^ intake in response to a voltage change (Fig. 2f and g). In some bacteria, the direction of flagellar rotation and chemotaxis changes depending on the internal Ca^2+^ concentration (Ordal, 1977; Tisa et al., 1993; Tisa and Adler, 1995). *M. ruber* was isolated from hot springs, and therefore a sufficient amount of Ca^2+^ is likely to exist in its native environment (Loginova et al., 1984). CavMr activation by a voltage change, which could vary depending on the environmental ionic conditions, might lead to any response to adapt to the new environment, such as flagellar rotation. These characteristics indicate the existence of signal coupling between the membrane voltage and Ca^2+^, even in the early stages of life, which might be the origin of the corresponding functions in eukaryotes, such as muscle contraction.

NavPp permeates more Na^+^ than Ca^2+^, but its selectivity is modest (Fig. 3f and Table 1). Notably, *P. pacifica* is a marine myxobacterium that requires NaCl for its growth (Iizuka et al., 2003). As mentioned above, the basic architecture of the CavMr/NavPp group is thought to produce a preference for Ca^2+^. *P. pacifica* might modify this channel architecture to acquire a Na^+^ intake pathway, which would likely result in the remaining feature of low-affinity Ca^2+^ inhibition in NavPp. This flexible usage of homo-tetrameric channels to allow different cations to permeate is also reported in another bacterium, *Bacillus alkalophilus* (DeCaen et al., 2014). NsvBa from *B. alkalophilus* is a non-selective channel whose selectivity filter is changed from “TLESWAS”, a typical Na^+^-selective sequence in alkaliphilic bacillus, to “TLDSWGS”, possibly to adapt to its ionic environment. Recently, an early eukaryote, diatom, was found to have another homo-tetrameric channel with no selectivity that has an important role in electrical signaling in this species (Helliwell et al., 2019). These findings suggest that the cation selectivity of the homo-tetrameric channel family can be flexibly tuned to realize the required roles specific to its original species. We also found four homologues that have filter sequences similar to CavMr “Tx(E/D)GWx(D/E)” in the NCBI database (WP_009945599.1, WP_012983075.1, WP_024079824.1, and XP_002186055.1). XP_002186055.1 belongs to diatoms, which implies the wide use of monomeric Cavs in various organisms – not only prokaryotes, but also eukaryotes.

### Insights into Ca^2+^ selectivity and the evolution of mammalian Cavs

Aspartate residues are generally observed in the Ca^2+^ permeation pathway in ion channels, as well as many Ca^2+^ binding proteins (Halling et al., 2016; Yan et al., 2015; Zalk et al., 2015). Actually, NavAb and NaChBac were successfully transformed to Ca^2+^-selective channels with the aspartate-introduced filter sequences “TLDDW(S/A)D” (Tang et al., 2014; Yue et al., 2002). But, our results elucidate that this strategy is not the only method for achieving high Ca^2+^ selectivity. Human Cavs subdomains possess, at most, two aspartate residues in the selectivity filters in other part than position 3. In addition, the negatively charged residue at position 3, which is thought to be the most critical for cation selectivity in both Navs and Cavs, is not aspartate, but glutamate in most of the human Cavs subdomains (Yu and Catterall, 2004). CavAb has 12 aspartates in the selectivity filter of a channel tetramer, while there are 4 aspartates in CavMr. The net negative charge is 5~7 in mammalian Cavs, 8 in CavMr, and 12 in CavAb. These findings indicate that Ca^2+^ selectivity can be achieved with even fewer negative charges than CavAb and close to mammalian Cav, probably with the contribution of a special backbone structure around the selectivity filters.

It is noteworthy that the selectivity filter sequence of CavMr is very similar to those of human Cav subdomains I and III, both of which possess a glycine at position 4 (Fig. 6c). Especially, the Cav3.1 and 3.2 subdomains I have the same sequence as CavMr. On the other hand, the sequences of subdomains II and IV are relatively similar to that of CavAb (Fig. 6d). These sequence similarities of the glycine residue at position 4 are also found in CatSper, the sperm calcium permeable channel (Darszon et al., 2011). The channel region of CatSper is formed by four different subunits (CatSper1-4). The selectivity filters of CatSper 3 “TVDGWTD” and CatSper 4 “TQDGWVD” are similar to that of CavMr and share a glycine residue at position 4 as well as subdomains I and III of 24TM Cavs. These similarities indicate the generality of the CavMr-like Ca^2+^-selectivity mechanism. Further investigation of the detailed structure of CavMr may help to elucidate the principles and origin underlying Ca^2+^ selectivity.

## Material and methods

### Cloning of BacNav homologues and site-directed mutagenesis

The NaChBac amino acid sequence (NP_242367) was used as the query for a BLASTP search against the Microbial Genomic database at NCBI. The identified primary sequence data were obtained from Entrez at NCBI (*Meiothermus ruber* as ZP_04038264, *Plesiocystis pacifica* as ZP_01909854, *Halomonas elongata* as YP_003896792 and *Teredinibacter turnerae* as YP_003073405). These DNAs were synthesized by Genscript Inc. and subcloned into the pCI vector (Promega) using the EcoRI and SalI sites and the pBiEX vector (Novagen) using NcoI and BamHI sites, respectively. Site-directed mutagenesis was achieved by polymerase chain reaction (PCR) of the full-length plasmid containing the Nav gene using PrimeSTAR^®^ MAX DNA Polymerase (Takara Bio.). All clones were confirmed by DNA sequencing.

### Electrophysiological analysis using mammalian cells

For the recordings related to mole fraction effects (Fig. S2a-d), currents were recorded from Chinese hamster ovary (CHO) -K1 cells expressing channels. The recordings were performed as described previously (Tateyama and Kubo, 2018). Cells were transfected with channel DNAs using the LipofectAMINE 2000 (Invitrogen) and plated onto cover slips. Currents were recorded 24-36 h after transfection. Current recording by the whole cell patch clamp technique was performed using Axopatch 200B amplifiers, Digidata1332A, and pClamp 9 software (Molecular Devices). The pipette solution contained 130 mM KCl, 5 mM Na_2_-ATP, 3 mM EGTA, 0.1 mM CaCl_2_, 4 mM MgCl_2_ and 10 mM HEPES (pH 7.2 adjusted with KOH). The bath solution contained 135 mM NaCl, 4 mM KCl, 1 mM CaCl_2_, 5 mM MgCl_2_ and 10 mM HEPES (pH 7.4 adjusted with NaOH). For the measurement of mole fraction effects, the bath solutions containing different ratio of NaCl / CaCl_2_ (135/0, 108/18, 81/36, 54/54, 27/82 and 0/90 mM) were used. The Ca^2+^-free solution was achieved with the solution containing 135 mM NaCl, 1 mM EGTA and 0 mM CaCl_2_.

### Electrophysiological measurement in insect cells

The recordings other than those for mole fraction effects were performed using SF-9 cells. SF-9 cells were grown in Sf-900™ III medium (Gibco) complemented with 0.5% 100× Antibiotic-Antimycotic (Gibco) at 27°C. Cells were transfected with target channel-cloned pBiEX vectors and enhanced green fluorescent protein (EGFP)-cloned pBiEX vectors using Fugene HD transfection reagent (Promega). The channel-cloned vector (2 μg) was mixed with 0.5 μg of the EGFP-cloned vector in 100μL of the culture medium. Next, 3 μL Fugene HD reagent was added and the mixture was incubated for 10 min before the transfection mixture was gently dropped onto cultured cells. After incubation for 16-48 h, the cells were used for electrophysiological measurements. In the measurement of I-V relation curves, the pipette solution contains 75 mM NaF, 40 mM CsF, 35 mM CsCl, 10 mM EGTA, and 10 mM HEPES (pH 7.4 adjusted by CsOH) For evaluation of ion selectivity, high Na pipette solution (115 mM NaF, 35 mM NaCl, 10 mM EGTA, and 10 mM HEPES (pH 7.4 adjusted by NaOH)) was used. For the evaluation of Ca, Sr, K and Cs selectivity, Ca solution (100 mM CaCl_2_, 10 mM HEPES (pH 7.4 adjusted by Ca(OH)_2_), and 10 mM glucose), Sr solution (100 mM SrCl_2_, 10 mM HEPES (pH 7.4 adjusted by Sr(OH)_2_), and 10 mM glucose), K solution (150 mM KCl, 2mM CaCl_2_, 10 mM HEPES (pH 7.4 adjusted by KOH), and 10 mM glucose), and Cs solution (150 mM CsCl, 2mM CaCl_2_, 10 mM HEPES (pH 7.4 adjusted by CsOH), and 10 mM glucose) were used as the bath solution, respectively. *E*_rev_ of high Ca^2+^ selective channels were measured under three external solutions containing 144mM NMDG-Cl and 4mM CaCl_2_, 135mM NMDG-Cl and 10mM CaCl_2_, 120mM NMDG-Cl and 20mM CaCl_2_ (10mM HEPES pH 7.4 adjusted with HCl). And *E*_rev_ of high Ca^2+^ selective channels for the calculation of *P*_K_/*P*_Ca_ and *P*_Cs_/*P*_Ca_ were measured under external solutions containing 135mM NMDG-Cl and 10mM CaCl_2_ (10mM HEPES pH 7.4 adjusted with HCl) with high K pipette solution (115 mM KF, 35 mM KCl, 10 mM EGTA, and 10 mM HEPES (pH 7.4 adjusted by KOH)) and high Cs pipette solution (115 mM CsF, 35 mM CsCl, 10 mM EGTA, and 10 mM HEPES (pH 7.4 adjusted by CsOH)), respectively. *E*_rev_ of NavPp for the calculation of *P*_Ca_/*P*_Na_ or *P*_Sr_/*P*_Na_ were measured under external solution containing 50mM NMDG-Cl, 40mM NaCl, 40mM CaCl_2_ or SrCl_2_ and 10mM HEPES pH 7.4 adjusted with NaOH. *E*_rev_ of NavPp for the calculation of *P*_Cs_/*P*_Na_ was measured under high Cs pipette solution and external solution containing 110mM NMDG-Cl, 40mM NaCl, 3mM CaCl_2_ and 10mM HEPES pH 7.4 adjusted with NaOH.

As the pipette solution for measurement of the Ca block effect in NavPp, low Na pipette solution (140 mM CsF, 10 mM NaCl, 10 mM EGTA, and 10 mM HEPES (pH 7.4 adjusted by CsOH)) and high Na pipette solution were used for inward and outward current measurement, respectively. As a bath solution, Ca blocking solution (30 mM NaCl, 120 mM NMDG-Cl, 1.5 mM CaCl_2_, 10 mM HEPES (pH 7.4 adjusted by NaOH) and 10 mM glucose) was used for the 1.5 mM Ca blocking condition. In 10mM Ca blocking condition, a bath solution contains 30 mM NaCl, 105 mM NMDG-Cl, 10 mM CaCl_2_, 10 mM HEPES (pH 7.4 adjusted by NaOH) and 10 mM glucose. And, in each Ca blocking conditions, 15 mM NMDG-Cl was replaced per 10 mM CaCl_2_. The bath solution was changed using the Dynaflow^®^ Resolve system. All experiments were conducted at 25 ± 2°C. All results are presented as mean ± standard error.

### Calculation of ion selectivity by the GHK equation

To determine the ion selectivity of each channel, the intracellular solution and extracellular solution were arbitrarily set and the reversal potential at each concentration was measured by giving the ramp pulse of membrane potential. The applied ramp pulse was set to include the reversal potential. In addition, a depolarization stimulus of 2-10 ms was inserted to check whether the behavior of the cell changed for each measurement. As a result, *P*_Ca_/*P*_Na_ was calculated by substituting the obtained reversal potential (*E*_rev_) into the expression derived from the GHK equation (Frazier et al., 2000);

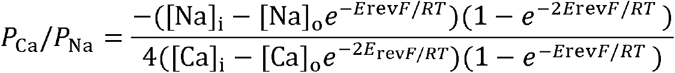

where *F* is Faraday’s constant, *R* is the Gas constant, and *T* is 298.1[K]. The same expression was used for Sr^2+^. The Sr^2+^ selectivity (*P*_Sr_/*P*_Na_) was measured in the same way.

Na^+^ selectivity against monovalent cations (*P*_M_/*P*_Na_) was calculated by substituting the obtained reversal potential and *P*_Ca_/*P*_Na_ into the expression derived from the GHK equation (Lopin et al., 2012):

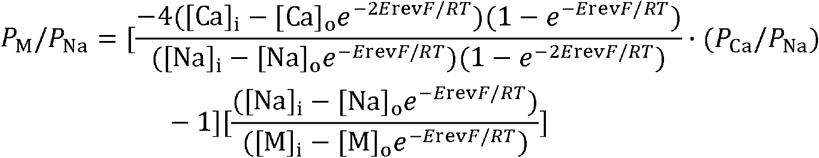

## Acknowledgements

We very appreciate to Dr. Yoshihiro Kubo for the critical suggestion about the structure of our manuscript. This work was supported by Grants-in-Aid for Scientific Research (S), a Grant-in-Aid for Young Scientists (B), the Japan Agency for Medical Research and Development and the Toyoaki Scholarship Foundation.

## Author contribution

T.S. and K.I. conducted the experiments; T.S. searched for homologues; Y.Y. and K.I. performed the electrophysiological experiments of insect cells; M.T. performed the electrophysiological experiments of mammalian cells; T.S., Y.Y., H.N. and K.I. optimized the measurement conditions; T.S., Y.Y., H.N., Y.F. and K.I. contributed to the study design and wrote the paper.

---

**Fig. S1.**
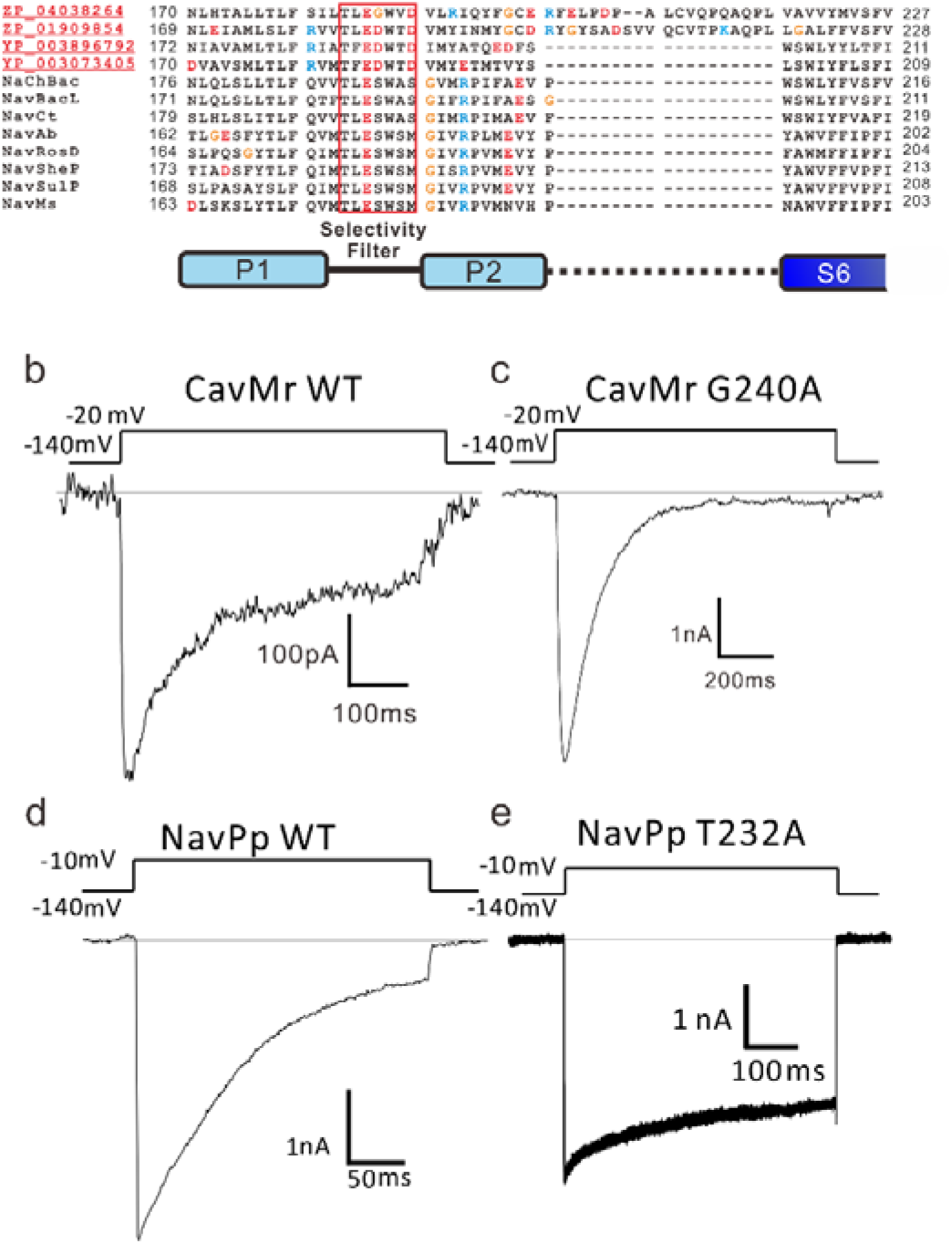
NavPp T232A and CavMr G240A mutants suppress inactivation. a). Alignment of the deduced amino acid sequences of P1 helix to P2 helix domain of novel cloned homologues with well characterized BacNavs. b) and c). Whole-cell current in CavMr wild type (WT; b) and a G240A mutant (c) when the pulse of −20 mV was given for 500 ms and 1sec, respectively. d) and e). Whole-cell recording of NavPp wild type (WT; d) and NavPp T232A mutant (e) when the pulse of −10 mV was given for 250 ms and 500ms, respectively.

**Fig. S2.**
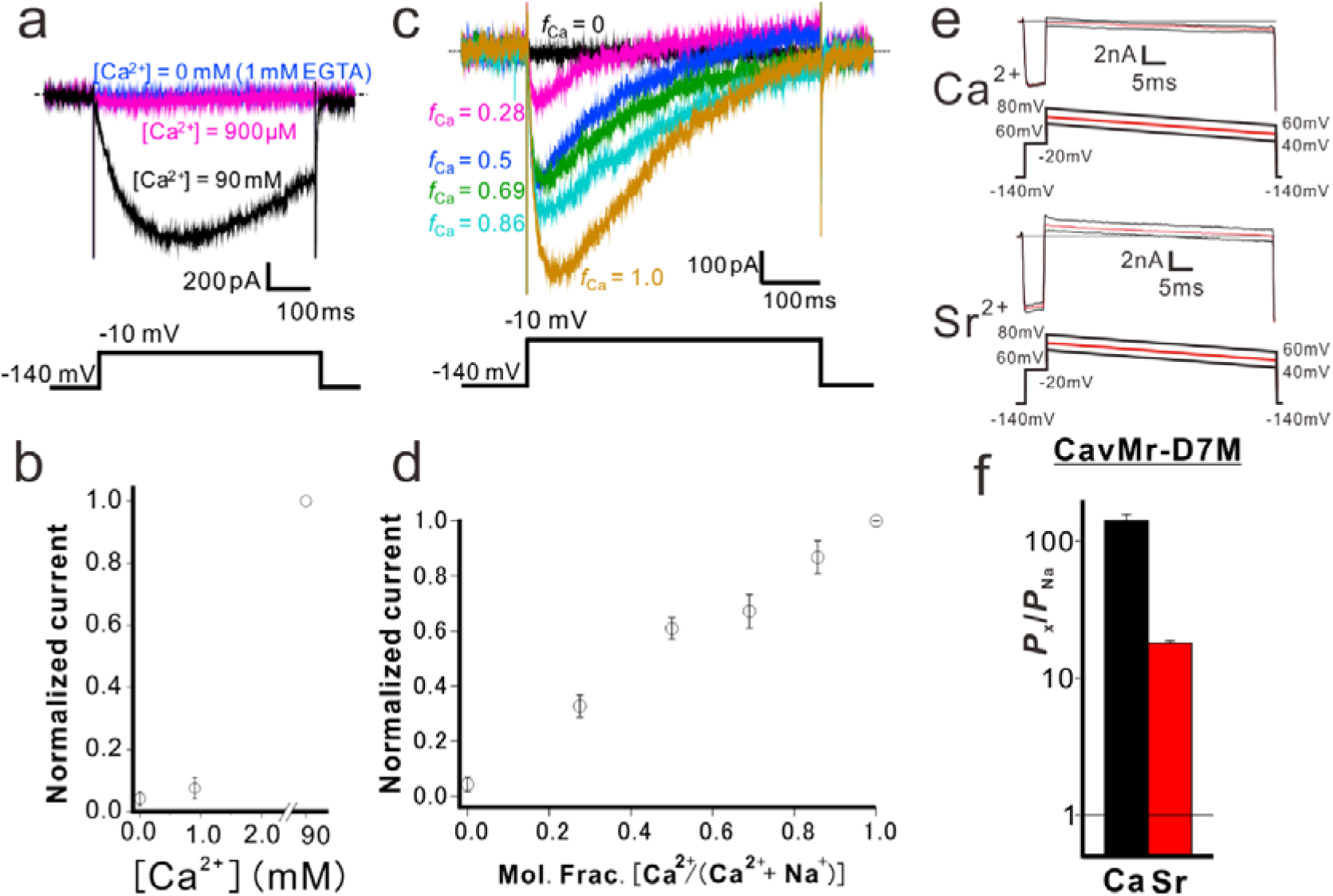
The characterization of the selectivity filter of CavMr. a). No observation of anomalous mole fraction effects in CavMr. CavMr currents were recorded in the bath solution containing the following concentration of Na^+^ and Ca^2+^; [Na^+^]: [Ca^2+^] is 0:90, 133.7:0.9, 135:0 (mM), respectively. The 0 mM Ca^2+^ solution also contains 1 mM EGTA. b). The plot of the normalized current amplitude of CavMr obtained from a. c). Representative current traces of CavMr under different mole fraction of Ca^2+^. *f*_Ca_ indicates [Ca^2+^]_out_ / ([Ca^2+^]_out_ + [Na^+^]_out_). d). The plot of the normalized current amplitude to each mole fraction, as measured in c. e). For the evaluation of the relative permeability of Ca^2+^ and Sr^2+^ to Na^+^ of CavMr-D7M, Ca solution (100mM CaCl_2_, 10mM HEPES (pH 7.4 adjusted with Ca(OH)_2_) and 10 mM glucose) and Sr solution (100 mM SrCl_2_ 10 mM HEPES (pH 7.4 adjusted by Sr(OH)_2_) and 10 mM glucose) were used as bath solution, respectively. High Na pipette solution (115 mM NaF, 35 mM NaCl, 10 mM EGTA, and 10 mM HEPES (pH 7.4 adjusted by NaOH)) was used. Currents were generated by the step pulse of −20 mV from −140 mV holding potential, followed by the ramp pulses with different voltage values. The time courses of the membrane potentials were shown at the bottom of each current traces. f) The relative permeability of divalent cation to Na^+^ in the CavMr-D7M, whose position 6 residue of the selectivity filter was neutralized by the corresponding residue of NavAb.

**Fig. S3.**
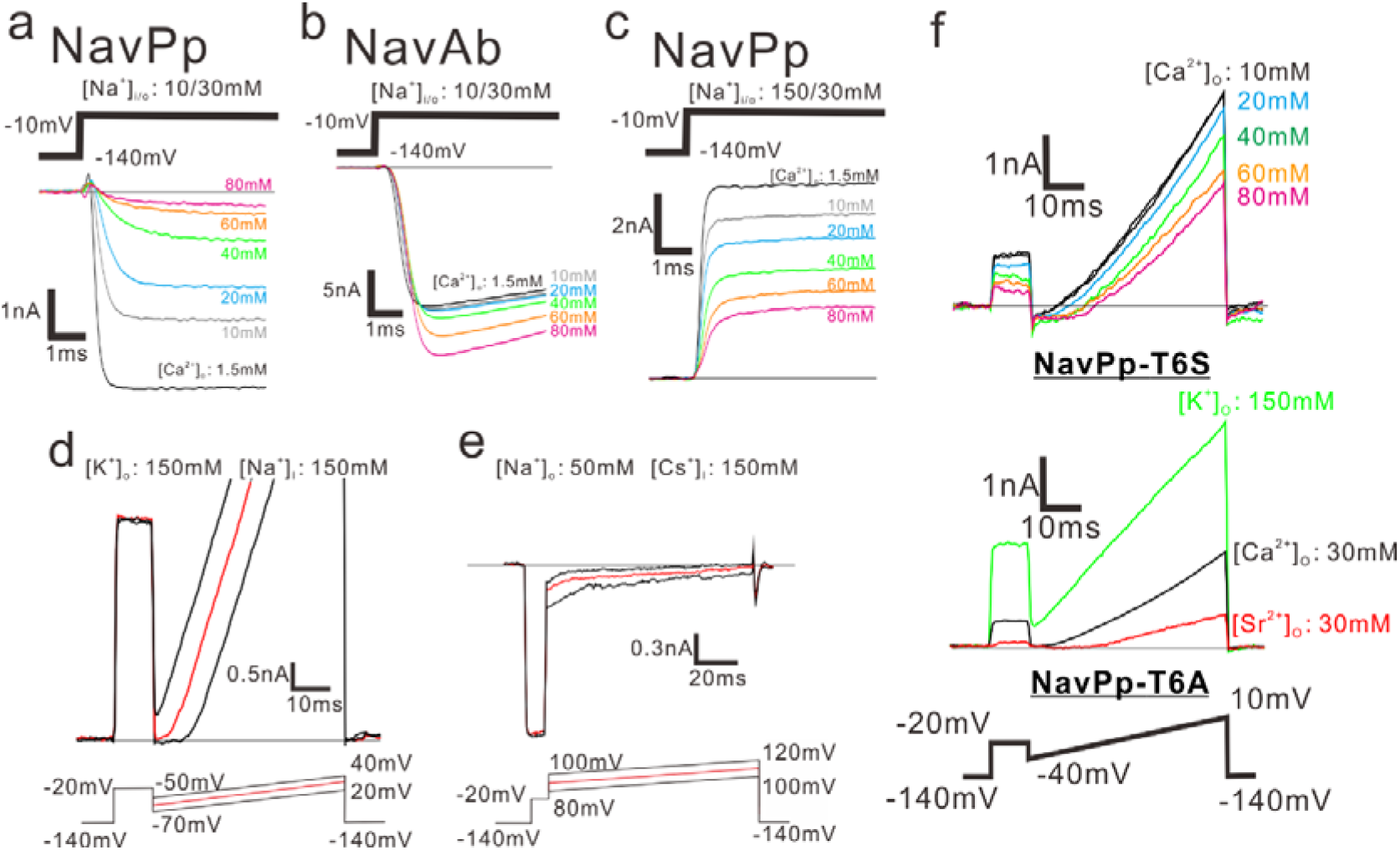
The characterization of the selectivity filter of NavPp. a) and b) NavPp and NavAb currents generated by the step pulses from −140 mV holding potential to −10 mV under the various [Ca^2+^]_out_ ranging from 1.5 to 80 mM. Inward currents are observed in the 30 mM [Na^+^]_out_ and 10 mM [Na^+^]_in_. c) NavPp currents generated by the same pulse protocol under various [Ca^2+^]_out_. Outward currents are observed in the 30 mM [Na^+^]_out_ and 150 mM [Na^+^]_in_. d and e) For the evaluation of the relative permeability of K^+^ and Cs^+^ to Na^+^ of NavPp, K solution (150 mM KCl, 2mM CaCl_2_ 10 mM HEPES (pH 7.4 adjusted by KOH) and 10 mM glucose) and low Na solution (50 mM NaCl, 100mM NMDG-HCl, 2mM CaCl_2_ 10 mM HEPES (pH 7.4 adjusted by NaOH) and 10 mM glucose) were used as bath solution, respectively. High Na pipette solution (115 mM NaF, 35 mM NaCl, 10 mM EGTA, and 10 mM HEPES (pH 7.4 adjusted by NaOH)) and Cs pipette solution (115 mM CsF, 35 mM CsCl, 10 mM EGTA, and 10 mM HEPES (pH 7.4 adjusted by CsOH)) were used for K^+^ and Cs^+^ selectivity, respectively. Currents were generated by the step pulse of −20 mV from −140 mV holding potential, followed by the ramp pulses with different voltage values. The time courses of the membrane potentials were shown at the bottom of each current traces. f) The relative permeability of each cation species to Na^+^ in the single-point mutants of NavPp. The selectivity filter of NavPp was changed to the Ca^2+^-selective canonical-BacNavs mutants (T6S; TLEDWSD and T6A; TLEDWAD).

**Fig. S4.**
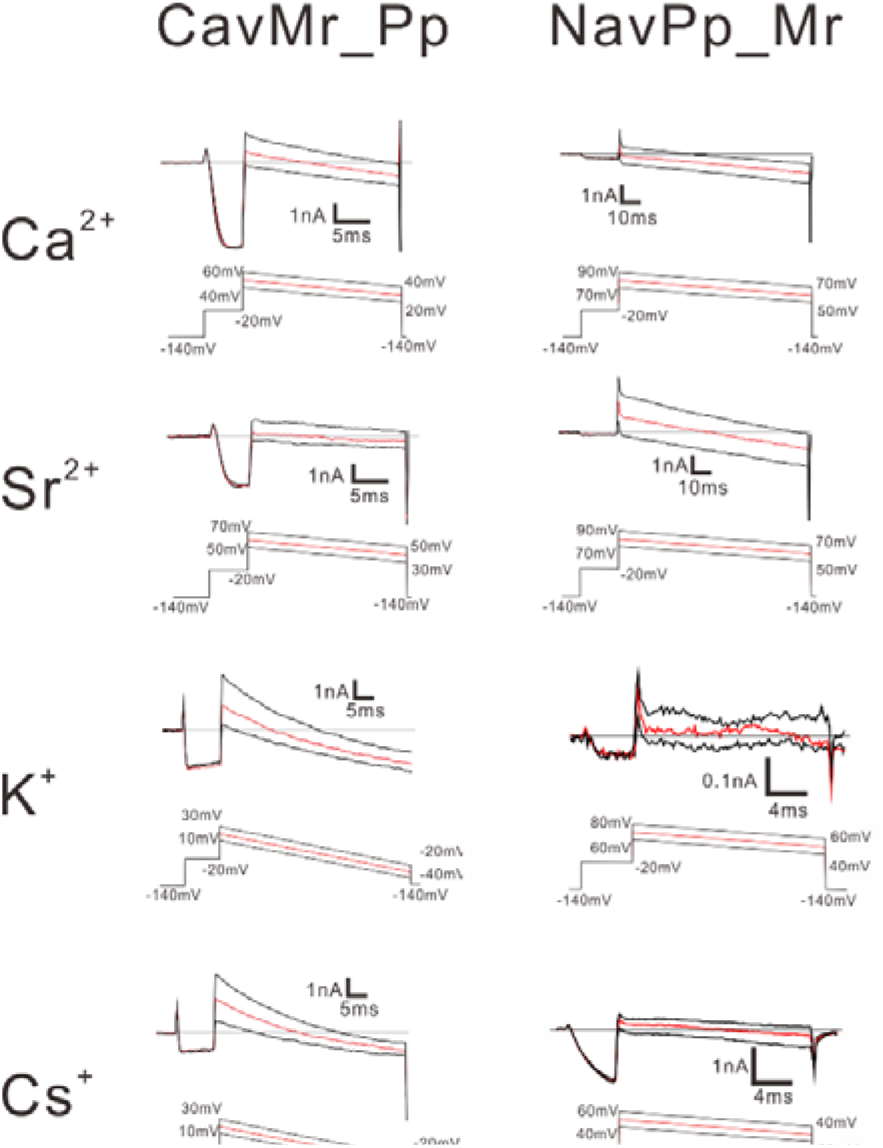
Representative current traces of the ramp pulse of selectivity filter-swapped mutants, CavMr_Pp and NavPp_Mr. For the evaluation of the relative permeability of Ca^2+^, Sr^2+^, K^+^ and Cs^+^ to Na^+^, Ca solution (100mM CaCl_2_, 10mM HEPES (pH 7.4 adjusted with Ca(OH)_2_) and 10 mM glucose), Sr solution (100 mM SrCl_2_, 10 mM HEPES (pH 7.4 adjusted by Sr(OH)_2_) and 10 mM glucose), K solution (150 mM KCl, 2mM CaCl_2_, 10 mM HEPES (pH 7.4 adjusted by KOH) and 10 mM glucose) and Cs solution (150 mM CsCl, 2mM CaCl_2_, 10 mM HEPES (pH 7.4 adjusted by CsOH) and 10 mM glucose) were used as bath solution, respectively. High Na pipette solution (115 mM NaF, 35 mM NaCl, 10 mM EGTA and 10 mM HEPES (pH 7.4 adjusted by NaOH)) was used. In the case of NavPp-Mr, the relative permeability of K^+^ and Cs^+^ to Ca^2+^ were evaluated in behalf of the relative permeability to Na^+^ because of high Ca selectivity of NavPp-Mr. For the evaluation, high K pipette solution (115 mM KF, 35 mM KCl, 10 mM EGTA and 10 mM HEPES (pH 7.4 adjusted by KOH)) and high Cs pipette solution (115 mM CsF, 35 mM CsCl, 10 mM EGTA and 10 mM HEPES (pH 7.4 adjusted by CsOH)) were used. As bath solution, Ca solution (100mM CaCl_2_, 10mM HEPES (pH 7.4 adjusted with Ca(OH)_2_) and 10 mM glucose was used. Currents were generated by the step pulse of −20 mV from −140 mV holding potential, followed by the ramp pulses with different voltage values. The time courses of the membrane potentials were shown at the bottom of each current traces.

**Fig. S5.**
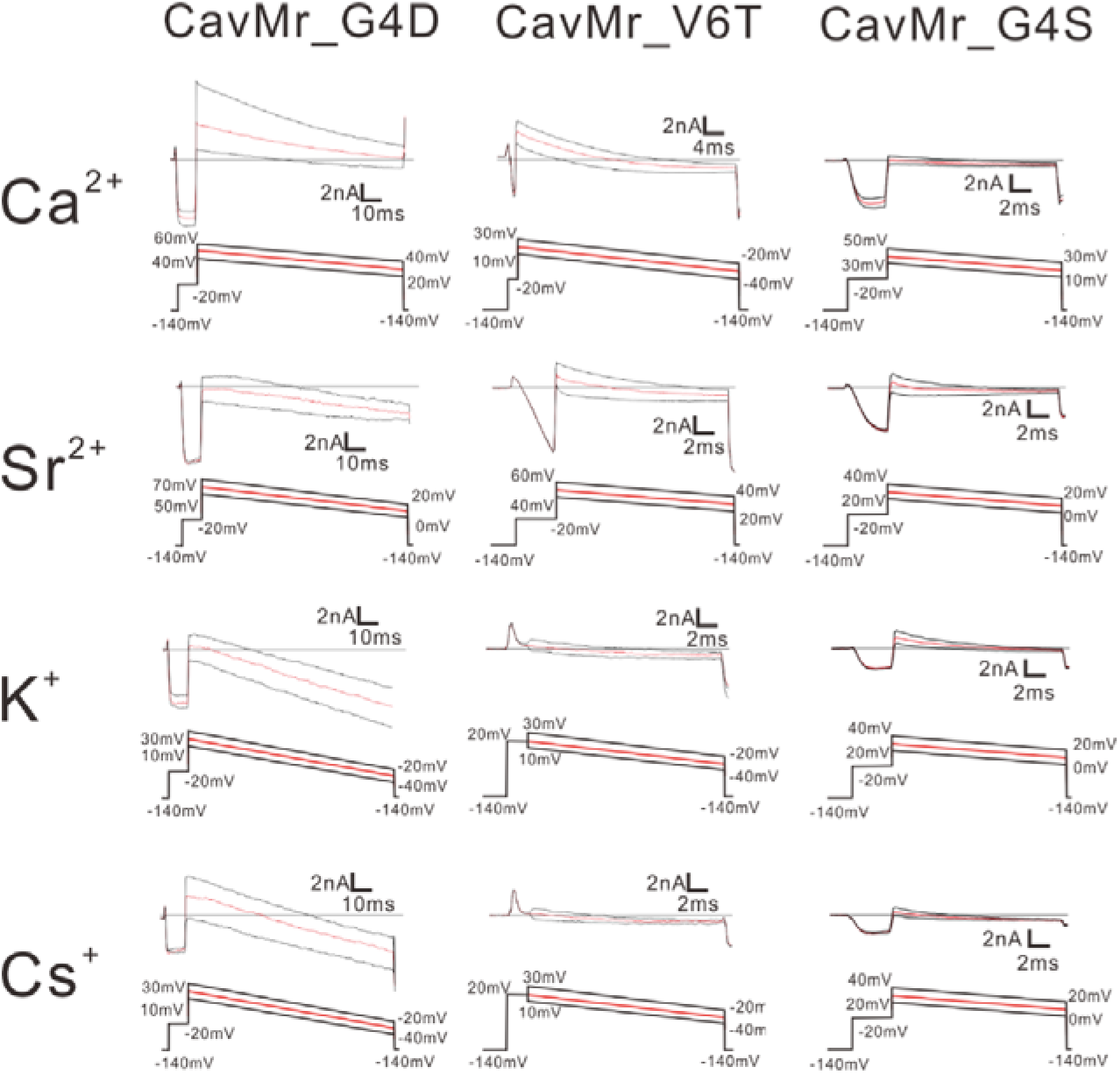
Representative current traces of the ramp pulse of single point-swapped mutants between CavMr. To evaluate the relative permeability of Ca^2+^, Sr^2+^, K^+^ and Cs^+^ to Na^+^, Ca solution, Sr solution, K solution and Cs solution were used as the bath solution, respectively. High Na pipette solution was used as the pipette solution. The solution contents were described in material method and Fig. S4. The time courses of the membrane potentials were shown at the bottom of each current traces.

**Fig. S6.**
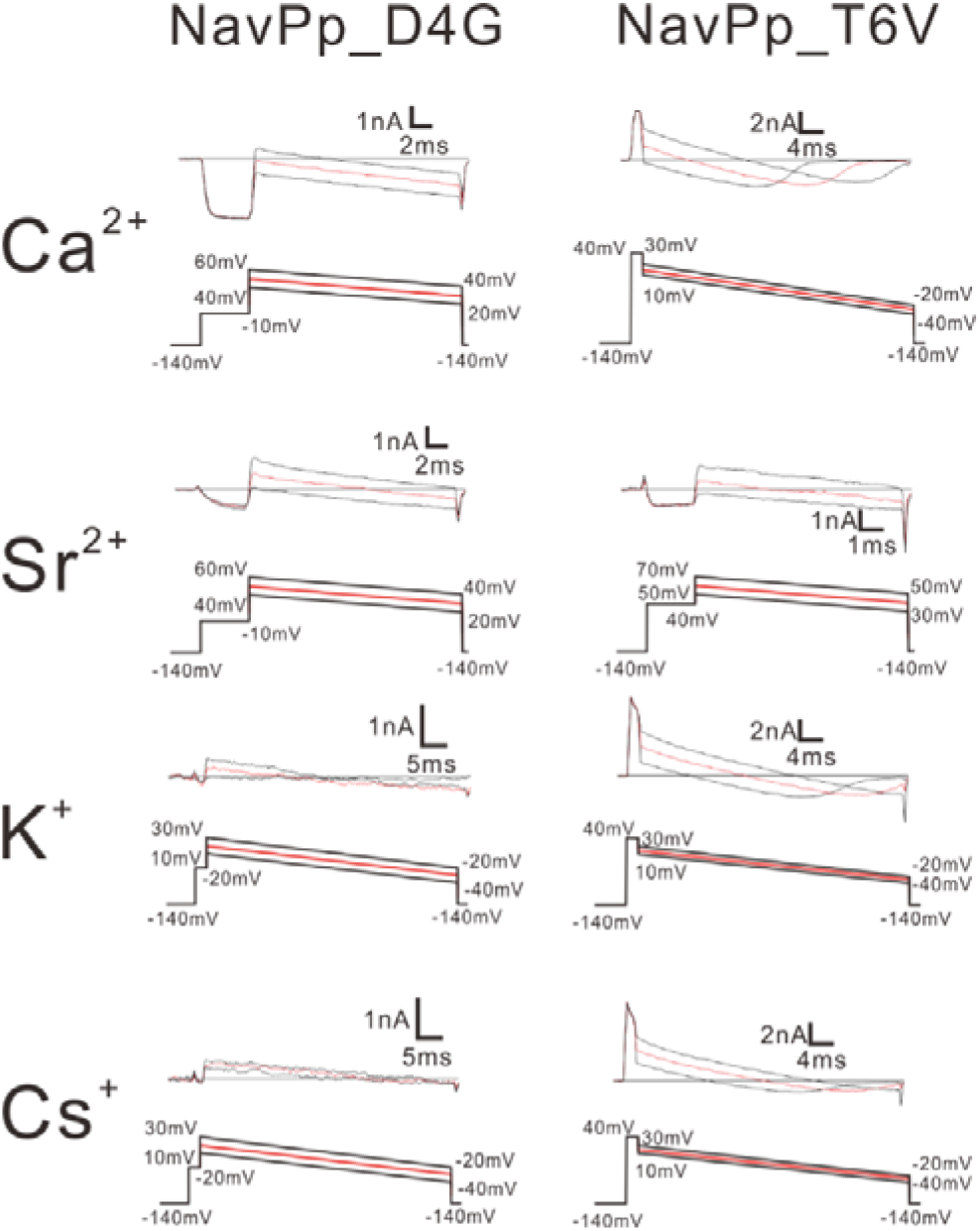
Representative current traces of the ramp pulse of single point-swapped mutants of NavPp. To evaluate the relative permeability of Ca^2+^, Sr^2+^, K^+^ and Cs^+^ to Na^+^, Ca solution, Sr solution, K solution and Cs solution were used as the bath solution, respectively. High Na pipette solution was used as the pipette solution. The solution contents were described in material method and Fig. S4. The time course of the membrane potentials was shown at the bottom of each current traces.

